## Supplemental Figures for "Gradient boosted decision trees reveal nuances of auditory discrimination behavior"

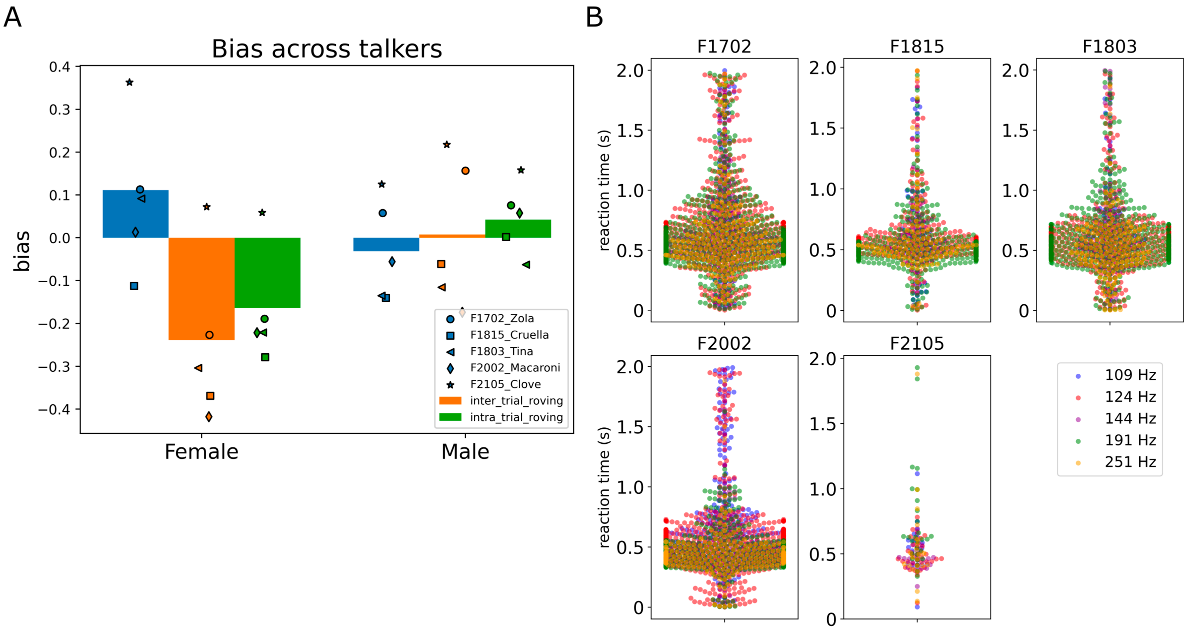


*Supplementary Figure 1. A, bias across trial conditions and talker types; B, reaction times of each animal for correct responses color-coded by F0 of the target word.*

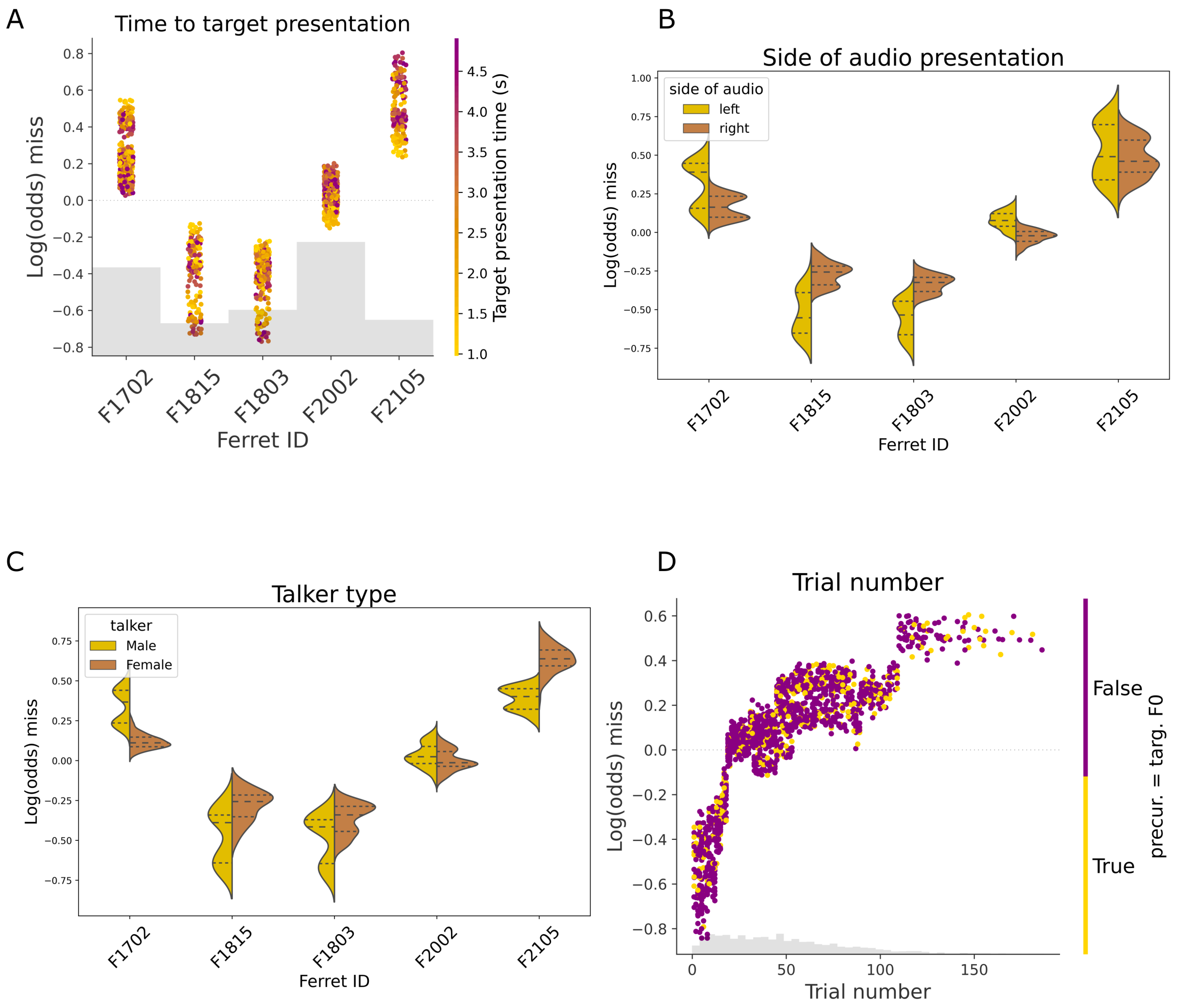


*Supplementary Figure 2. Partial dependency plots for the correct hit response/miss response model. A; SHAP values over the ferret ID color-coded by target presentation time; B, SHAP values over ferret ID color-coded by the side of audio presentation; C, same as B but color-coded by talker type; D, SHAP values over trial number color-coded by whether the trial had the precursor word F0 equal to the target F0.*

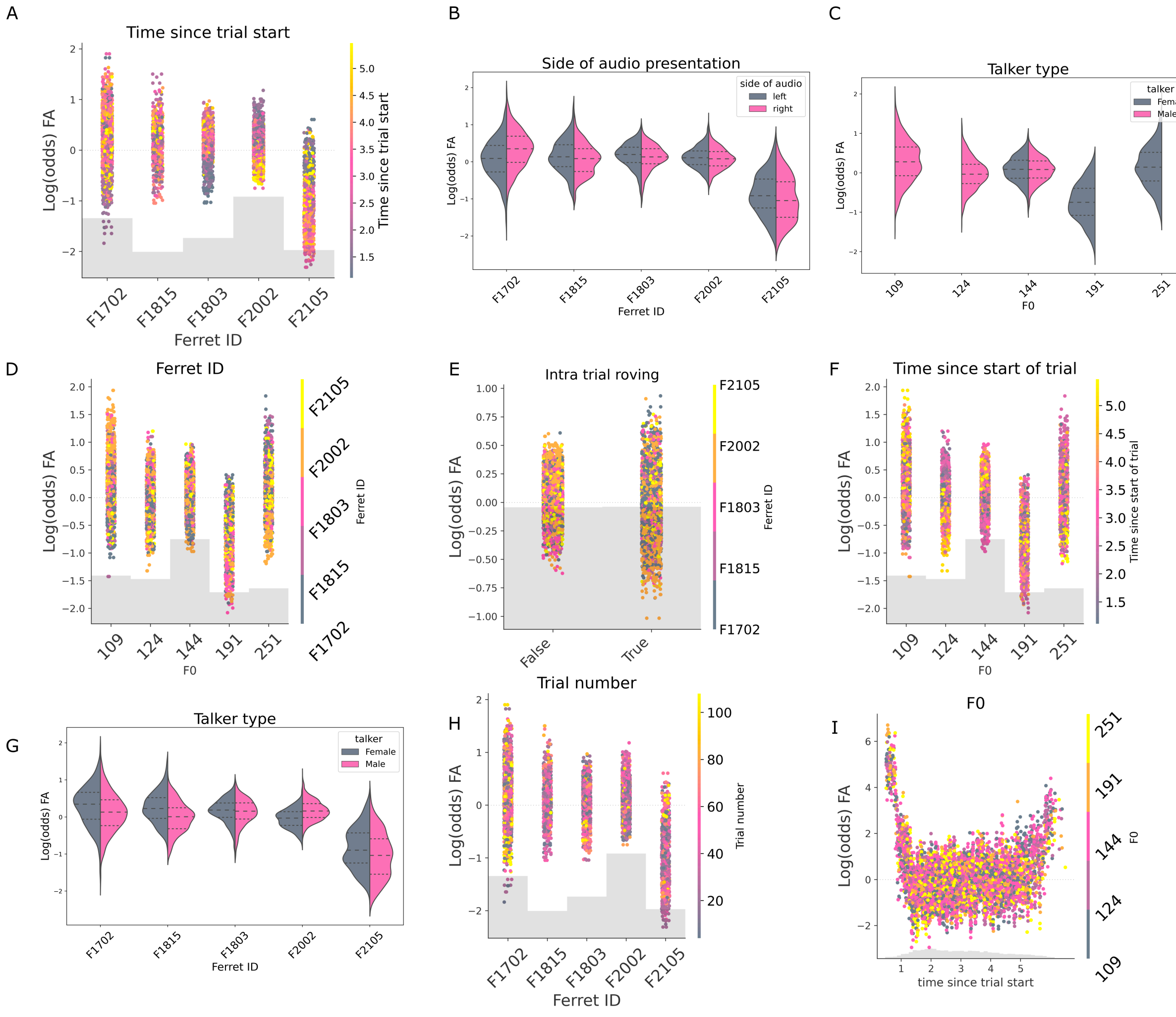


*Supplementary Figure 3. Partial dependency plots for the correct reject/false alarm model. A, partial dependency plot depicting the mean SHAP impact over the ferret ID color-coded by time within the trial; B, violin plot of the SHAP value over the ferret ID color-coded by the side of audio presentation; C, violin plot of the SHAP values over the F0 of the trial color-coded by talker type; D; SHAP partial dependency plots of false alarm likelihood by F0, color-coded by ferret ID; E, SHAP values over the F0 of the stream color-coded by trial number; F, same as E but color-coded by time since the start of the trial; Note that while the 191Hz F0 is associated with a higher false alarm rate, this should be interpreted in the context of the much lower FA rate associated with the female talker. G, violin plot of the SHAP value over ferret ID color-coded by talker type; H, SHAP value over ferret ID color-coded by trial number; I, SHAP*

*value over trial duration color-coded by F0.*

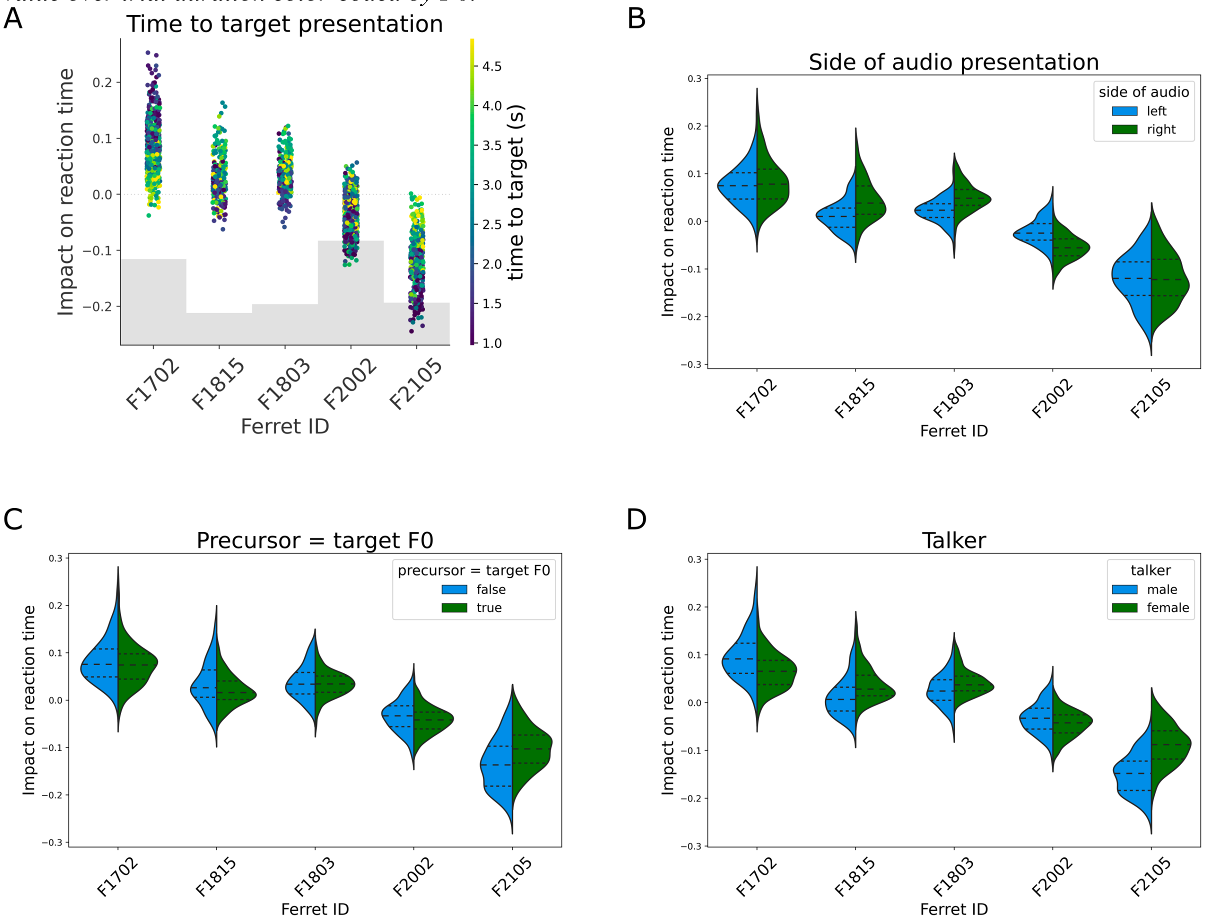


*Supplementary Figure 4. Correct hit response reaction time model partial dependency plots. A, SHAP values over the ferret ID color-coded by the time to target presentation; B, violin plot of the SHAP value over ferret ID color-coded by the side of audio presentation; C, same as B but color-coded by whether the precursor word's F0 was the same as the target word's F0; D, same as C but color-coded by the talker type for the trial.*

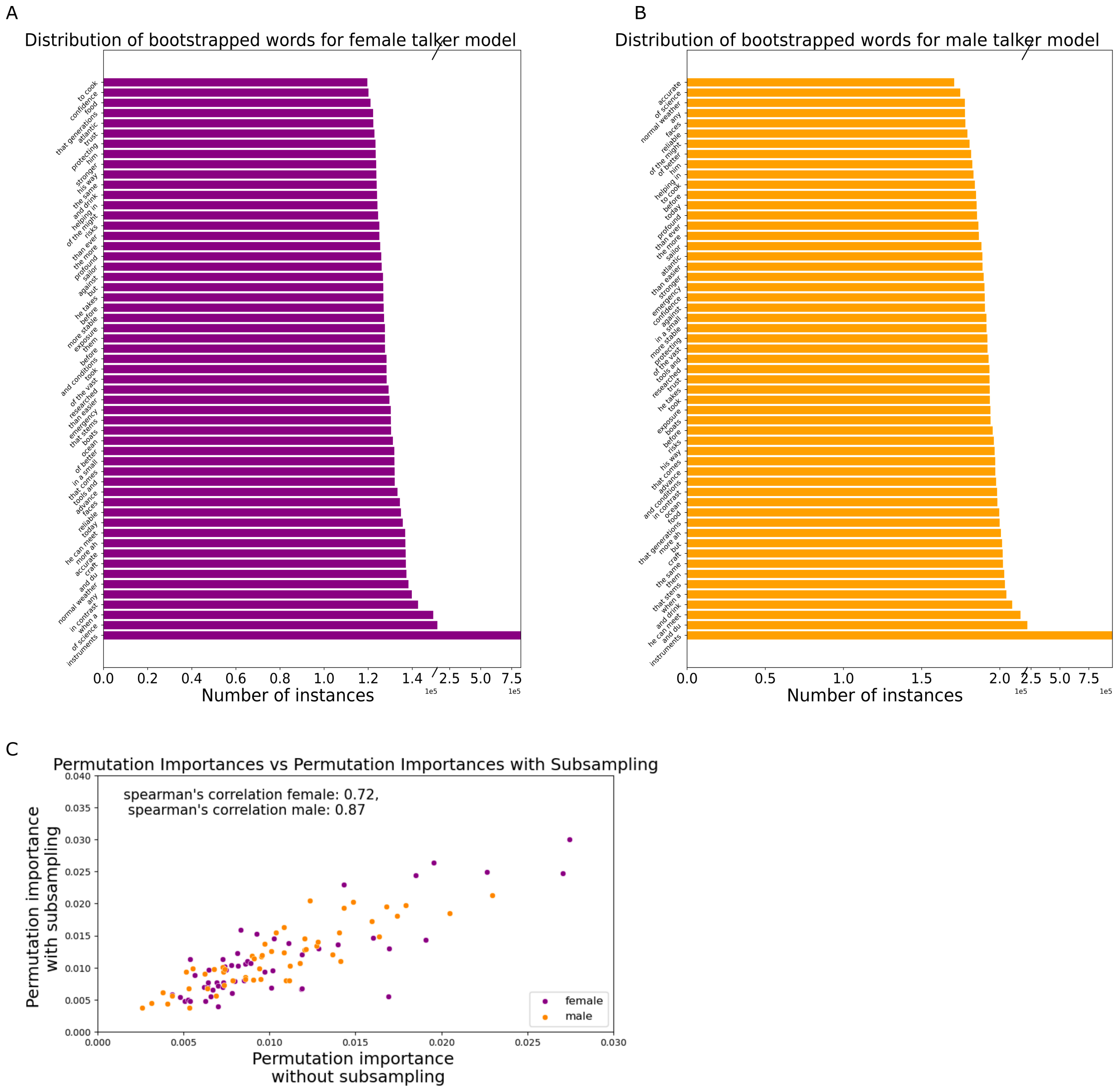


*Supplementary Figure 5. A, distribution of the probability of occurrence in the resampled dataset used for the response time model in Figure 5; B, same as A but for the male talker absolute reaction time model; C, scatter plot of the permutation importance of each word with subsampling to equalize the frequency to the distribution plotted in A and B, versus the corresponding permutation importance scores obtained from a model with the uncorrected word distributions.*

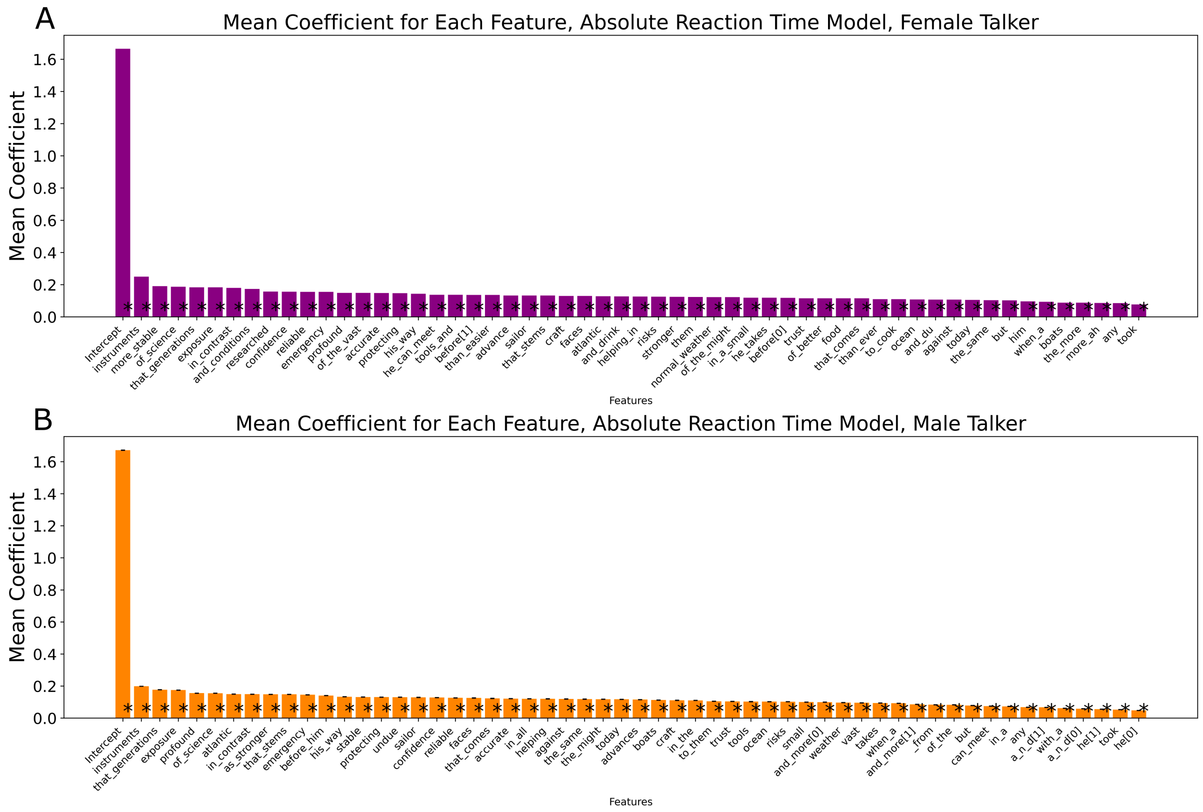


*Supplementary Figure 6. Average coefficients for the mixed effects model predicting the absolute reaction time of A, the female talker and B, the male talker. Asterisks represent mean p-values < 0.05. Error bars represent standard deviation.*
